# Genomic signatures associated with convergent funnel-web building behavior in spiders

**DOI:** 10.1101/2025.10.22.684048

**Authors:** Zheng Fan, Lu-Yu Wang, Zhi Li, Tian-Yu Ren, Bing Tan, Wei Pu, Wen-Hui Wu, Jun-Han Xiong, Ling-Xin Cheng, Jin-Xia Kong, Bin Luo, Zi-Zhong Yang, Chao Tong, Zhi-Sheng Zhang

## Abstract

Spiders exhibited diverse and intricate web-building behaviors, which represent a classic model for studying the evolution of complex traits. The funnel-web, a sheet-like web with a retreat tube, has evolved independently in several spider lineages, especially in the families Agelenidae and Macrothelidae. This repeated emergence of a complex behavior offers an example of this funnel-web building behavioral convergence. Here, we present a chromosome-level genome assembly of the agelenid spider *Tamgrinia laticeps*, along with annotated genomes of another agelenid spider *Eratigena atrica*, and a macrothelid *Orientothele yani* which convergently evolved funnel-web building behavior. Comparative genomic analysis of 15 spider species with diverse web-types revealed convergent signatures associated with funnel-web construction. Hundreds of genes tended to experience convergent shift in selective pressure, convergent positive section, and harbored convergent amino acid substitutions. These genes are associated with synaptic transmission (e.g., *SLC6A3, Gabbr1, SNAP25*), neuromuscular coordination (e.g., *ine, VACh*T), and behavioral regulation (e.g., *Crtc1, Lrrc7*). Notably, we detected identical convergent amino acid changes in *VAChT* and *ine* across all three funnel-web builders. Our findings demonstrate that convergent evolution of web architecture is linked to nervous system evolution, providing genomic insights into the basis of behavioral convergence.

## Introduction

Spiders exhibit an extraordinary diversity of web architectures, with the four best-known types being orb webs, sheet webs, tangle webs and funnel webs (Petruzzello et al., 2025; Foelix, 2011). Each type represents a distinct strategy for catching prey. These silk constructions not only serve as efficient tools for prey capture and shelter but also as striking examples of behavioral complexity in invertebrates. Funnel-web spiders typically inhabit cool, moist, and sheltered environments. Their webs consist of dense, horizontal sheets with a tubular retreat, in contrast to the vertical and circular orb webs. Designed for ambush rather than aerial prey capture, funnel webs are easily distinguished from those of orb-weaving or cobweb spiders. The spider remains concealed within the retreat during the day and waits near the entrance at night. When an insect or small animal disturbs the trip lines, the spider rapidly emerges, subdues the prey, and drags it back into the funnel. This ambush strategy is highly efficient and well adapted to nocturnal hunting (Nentwig et al., 2024). The instances of independent funnel-web building behaviors in distantly related lineages including Agelenidae and Macrothelidae provides an ideal system for studying the molecular basis of convergent behavioral evolution.

Recent studies have begun to uncover the neural and genetic basis of web-building behavior in spiders. Single-cell transcriptomic analyses in a recent study have revealed that neuronal diversification and molecular adaptation in neural genes underpin the emergence of complex web-building behaviors^4^ (Jin et al., 2024). Comparative neuroanatomical studies further demonstrate that web-building spiders possess distinct brain architectures compared with cursorial hunters, including reductions in visual neuropils and mushroom bodies but enlarged mechanosensory regions (Lehmann et al., 2023). These findings suggest that evolutionary transitions in web-building strategies are closely linked to reorganization of neural circuits and sensory processing. Environmental and physiological factors can also affect spider neurobiology and web construction. For instance, artificial light at night (ALAN) has been shown to induce measurable reductions in brain volume and potential neural damage in orb-weaving spiders (*Hortophora biapicata*), likely due to oxidative stress (Garwood et al., 2024). Similarly, aging has been associated with decreased brain volume and increased anomalies in web geometry, implying that changes in neural investment directly influence web-building performance^7^ (Pasquet et al., 2017). Moreover, neurophysiological studies on Australian funnel-web spiders revealed that venom-derived peptide toxins, such as N-atracotoxins and versutoxin, strongly modulate synaptic transmission in mammalian and insect neurons (Fletcher et al., 1997; Nicholson et al., 2004), highlighting the evolutionary refinement of neuroactive molecules in this lineage. Together, these studies indicate that both intrinsic (genetic and neural) and extrinsic (environmental) factors shape the evolution and function of the spider nervous system in web-building behaviors.

However, despite increasing evidence linking neural evolution to behavioral innovation, the genomic signatures associated with convergent evolution of web-building behavior across different spider lineages remain largely unknown. Here, we investigate the genomic basis of convergent funnel-web building behavior in 15 spider species representing at least three independent origins of funel-web building behaviors. We integrated comparative genomics and molecular evolution analyses to identify convergence at genome-wide, genic, and substitution site levels across spiders.

## Results

### Genome features of *Tamgrinia laticeps, Orientothele yani* and *Eratigena atrica*

In this study, we generated a high-quality, chromosome-level genome assembly of *T. laticeps*, with a total length of 7.38 Gbp and a scaffold N50 of 356.28 Mbp across 20 chromosomes. BUSCO assessment identified 94.5% of the conserved Arachnida_odb10 genes, indicating high assembly completeness. Repetitive elements accounted for approximately 66.72% of the genome (Figure 1C), dominated by DNA transposons (15.68%), LTR retrotransposons (9.70%), LINEs (2.92%), rolling-circle elements (0.64%), SINEs (0.07%), and unclassified repeats (38.35%). A total of 28,194 protein-coding genes were predicted and functionally annotated, with a BUSCO completeness score of 96.1%. We also re-annotated the genome of *Eratigena atrica*, in which interspersed repeats accounted for 49.91% of the genome (Figure 1C), including 0.21% SINEs, 3.96% LINEs, 2.40% LTR elements, 9.76% DNA transposons, 0.04% Rolling-circles and 33.58% unclassified repeats. A total of 47,673 protein-coding genes were predicted and functionally annotated, with a BUSCO completeness score of 91.8%.

**Figure 1.**
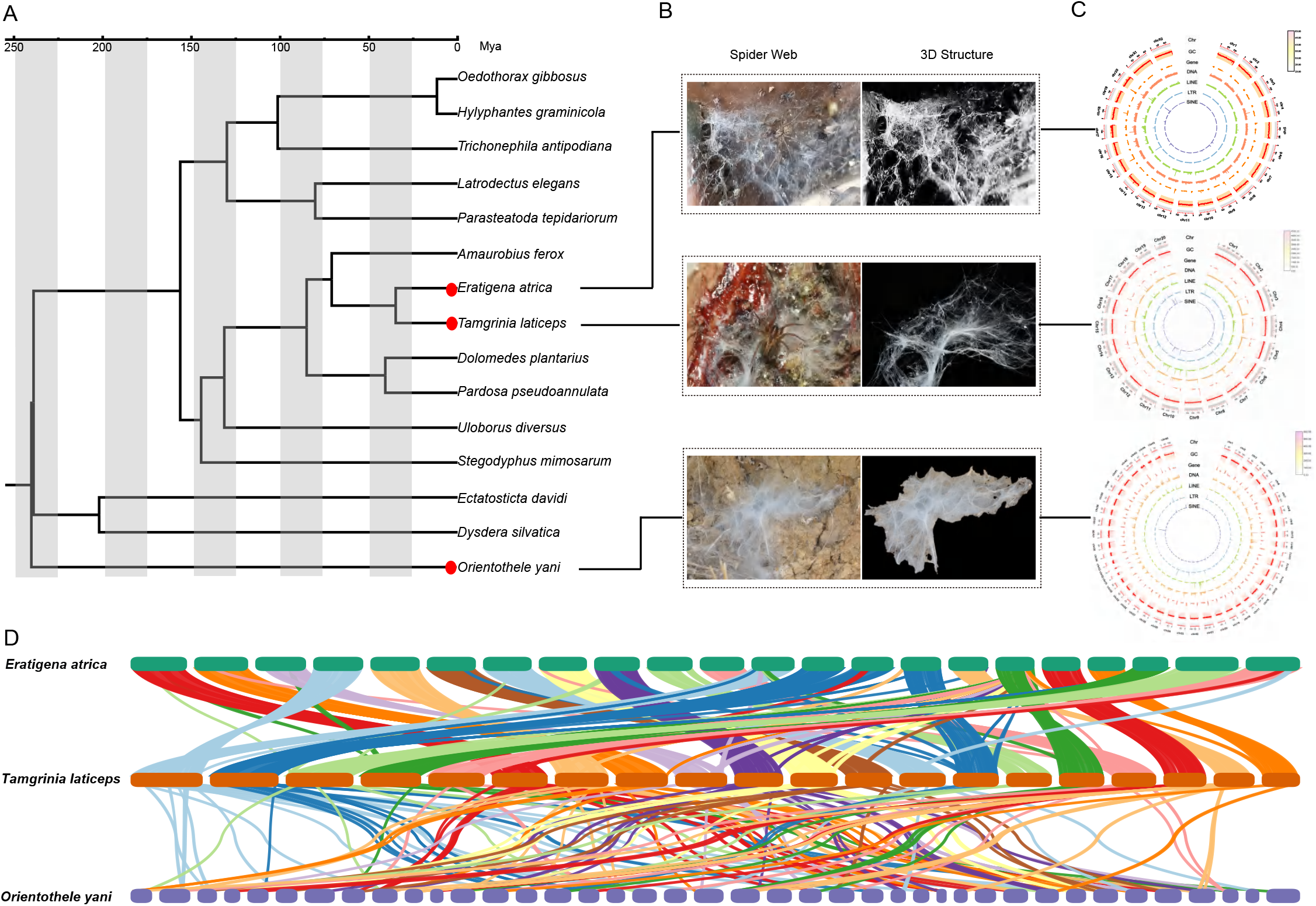
Phylogenetic relationship, web structure, genome characteristic, and collinearity analysis of funnel-web building spiders. (A) A phylogenetic tree of 15 spider species, including the three funnel-web weavers *Tamgrinia laticeps, Eratigena atrica*, and *Orientothele yani*, which are highlighted with red dots. (B) Photographs and corresponding 3D structural models of the webs constructed by *T. laticeps, E. atrica*, and *O. yani*. (C) Genome structure analysis in *T. laticeps, E. atrica*, and *O. yani*. (D) Syntenic relationships within the genomes of *T. laticeps, E. atrica*, and *O. yani*.

For *Orientothele yani*, interspersed repeats constituted approximately 67.81% of the genome (Figure 1C), comprising 0.74% SINEs, 14.45% LINEs, 1.03% LTR elements, 25.50% DNA transposons, 0.95% Rolling-circles and 26.09% unclassified repeats. A total of 24,173 protein-coding genes were predicted and annotated, achieving a BUSCO completeness of 88.7%.

### Phylogenetic relationships and convergent evolution of funnel-web architecture

To ensure reliable phylogenetic inference, we reconstructed a phylogenetic tree using 2,915 complete single-copy orthologs retrieved from the Arachnida_odb10 BUSCO dataset (n=2,934) among major spider lineages with high-quality genomes (Figure 1A). The results revealed that *T. laticep*s (Agelenidae) and *E. atrica* (Agelenidae) form a well-supported clade within the family Agelenidae, while *O. yani* belongs to the distantly related family Macrothelidae. Despite their large phylogenetic distance, all three species independently evolved a characteristic funnel-shaped web. Morphological observations and three-dimensional web models further confirmed the strong convergence in web architecture among these taxa (Figure 1B).

Comparative genomic analyses revealed some differences in chromosomal among the three funnel-web-building spiders (Figure 1D). *T. laticeps* and *E. atrica* (Agelenidae) exhibited high levels of chromosomal synteny and similar karyotypic structures, containing 20 and 22 chromosomes, respectively, consistent with their relatively recent divergence around 34.9 million years ago. In contrast, *O. yani* (Macrothelidae), which diverged from Agelenidae approximately 238.97 million years ago, possessed a highly distinct karyotype with 46 chromosomes and displayed extensive genome rearrangements and disrupted synteny relative to both Agelenid species. These findings indicate that large-scale genome organization remains relatively conserved within Agelenidae but has been substantially reorganized in *O. yani* since its early divergence. Remarkably, despite these deep genomic and evolutionary separations, all three species independently evolved morphologically similar funnel-web architectures, suggesting that convergent behavioral evolution can arise from distinct genomic frameworks shaped by long-term divergence.

### Evolutionary rate shifts associated with funnel-web evolution

To investigate molecular signatures associated with funnel-web evolution, we estimated relative evolutionary rates (RERs) for 8,765 shared orthologous, which reflect whether a given gene is evolving faster or slower than expected along specific branches of the phylogeny. Using this approach, we identified 212 rapidly evolving genes (Rho > 0, q-value < 0.05) and 292 slowly evolving genes (Rho < 0, q-value < 0.05) in funnel-web spiders (Figure 2A).

**Figure 2.**
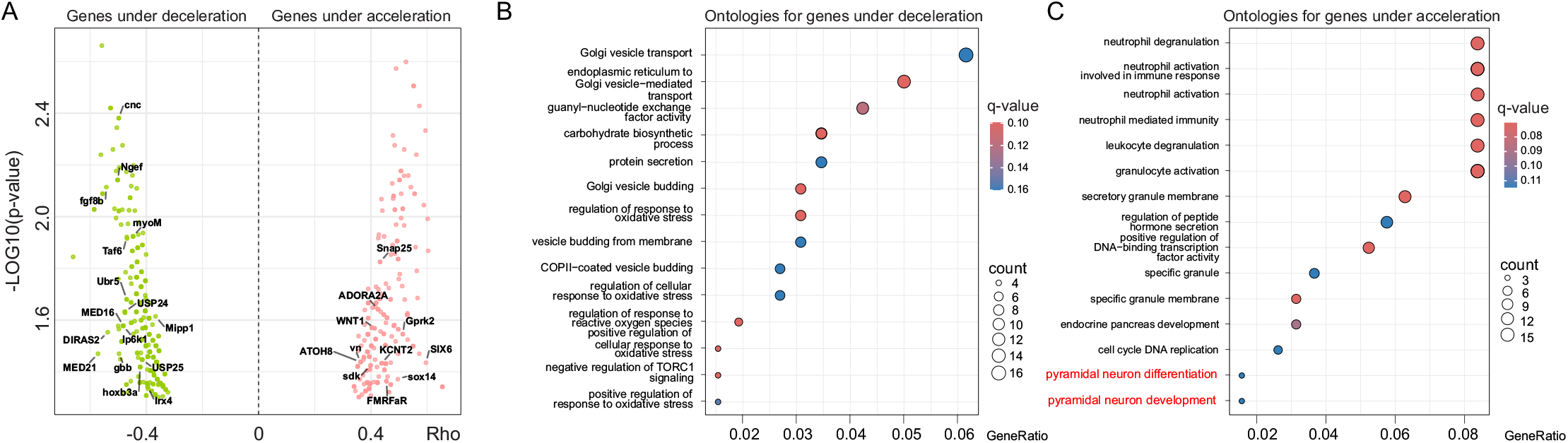
Selection pressure and functional analysis of genes in funnel-web building spiders. (A) Volcano plot depicting genes under significant deceleration (stabilizing selection) and acceleration (positive selection) in funnel-web weavers compared to other spiders. (B) Gene Ontology (GO) enrichment analysis for genes under significant deceleration. (C) Gene Ontology (GO) enrichment analysis for genes under significant acceleration.

Functional annotation of these genes revealed a subset potentially involved in neural regulation and behavioral control. Notably, candidate neural-related genes among the rapidly evolving set included *SLC6A3, Gabbr1, Snap25, Kcnab2*, and *ADORA2A*, whereas neural-related slowly evolving genes included *rpl27a, STX5, NDUFA2*, and *Irx4* (Figure 2A, Table S3).

GO enrichment analyses further highlighted distinct functional patterns (Figure 2B). Rapidly evolving genes were enriched for categories related to neural development and signaling, including pyramidal neuron differentiation and development, regulation of peptide hormone secretion, and positive regulation of DNA-binding transcription factor activity. In contrast, slowly evolving genes were enriched for processes associated with cellular homeostasis and protein trafficking, such as Golgi vesicle transport, protein secretion, and regulation of response to oxidative stress.

These results suggest that accelerated evolution of neural-related genes may contribute to behavioral innovation, facilitating the convergent evolution of funnel-web building, while conserved slowly evolving genes maintain essential cellular functions, providing a stable genomic and cellular background for the emergence of complex web-building behavior across divergent spider lineages.

### Relaxed and intensified selection in neural regulatory genes

To investigate selective pressures on neural regulatory genes in funnel-web spiders, we applied the RELAX method to assess gene-wide relaxation or intensification of selection(Figure 3A). Among the 575 genes showing relaxed selection (K < 1, q-value < 0.05), *FASN* (fatty acid synthase; K = 0.6957, q-value = 0.0011) exemplifies strong relaxation (Figure 3B, Table S4). *FASN* plays a central role in lipid biosynthesis and energy metabolism, processes that are essential for maintaining neuronal membrane composition and signaling. Its relaxed selection in funnel-web spiders may indicate reduced evolutionary constraint, potentially allowing metabolic flexibility to support energetically demanding behaviors such as web construction. Conversely, among the 100 genes under intensified selection (K > 1, q-value < 0.05), *Syt6* (synaptotagmin-6; K = 23.3, q-value = 0.0156) exhibits strong intensified selection (Figure 3B). *Syt6* is a key regulator of calcium-dependent neurotransmitter release at synapses, and its strong selective pressure suggests enhanced functional constraint in funnel-web spiders, likely supporting precise synaptic transmission required for the complex motor coordination during funnel-web building.

**Figure 3.**
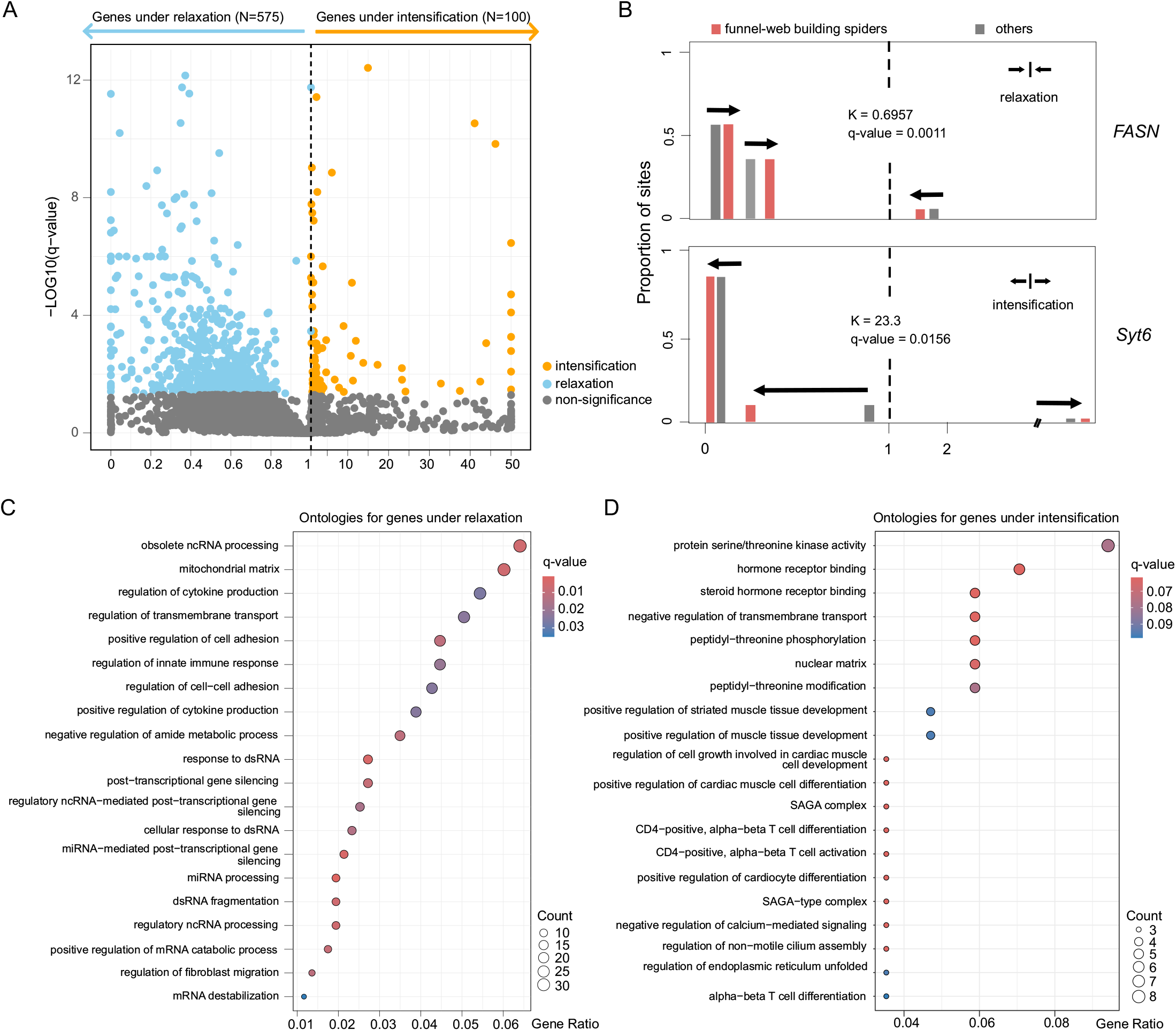
Selective pressure shift and functional profiling of genes in funnel-web building spiders. (A) Volcano plot illustrating genes that have undergone significant relaxation (weakened constraint) and intensification (strengthened constraint) of selective pressure in funnel-web builders compared to other spiders. (B) Detailed view of representative genes under relaxation (FASN) and intensification (Syt6), showing their evolutionary dynamics between funnel-web builders and other spiders. (C) Gene Ontology (GO) terms significantly enriched for genes under relaxed selection. (D) Gene Ontology (GO) terms significantly enriched for genes under intensified selection.

GO enrichment analysis revealed that relaxed selection genes were enriched in RNA processing, post-transcriptional regulation, immune response, and cell adhesion pathways, including obsolete ncRNA processing, mitochondrial matrix, regulation of cytokine production, and positive regulation of mRNA catabolic process (Figure 3C). These results suggest that reduced selective constraints may allow greater flexibility in regulatory networks, potentially facilitating behavioral innovation. In contrast, genes under intensified selection were enriched for kinase activity, hormone receptor binding, muscle and neural development, and transcriptional regulation, including protein serine/threonine kinase activity, steroid hormone receptor binding, positive regulation of muscle tissue development, and SAGA complex (Figure 3D). The elevated selective pressure on these genes indicates that precise neuromuscular and neural functions are maintained, likely supporting the complex motor and behavioral demands of funnel-web construction.

Overall, the RELAX analysis highlights a complementary pattern of selective pressures: relaxed selection permits flexibility in metabolic and regulatory pathways, whereas intensified selection preserves neural and synaptic function, together shaping the molecular architecture underlying the evolution of funnel-web building behavior.

### Positive selection in neural and regulatory genes

Genome-wide positive selection analysis revealed a set of genes showing adaptive evolution in funnel-web building spiders compared with other web types. A total of 102 genes were identified as positively selected (q-value < 0.05), several neural regulatory genes showed strong signals of positive selection, including *SLC6A3, Gabbr1, Crtc1, Lrrc7* (Densin-180), *Pak3, DCLK1*, and *NFIX* (Figure 4A, Table S5). These genes are known to modulate synaptic plasticity, learning, and memory, implying that adaptive evolution in neural circuits may underlie the refinement of web-building and prey-capture behaviors in funnel-web spiders.

**Figure 4.**
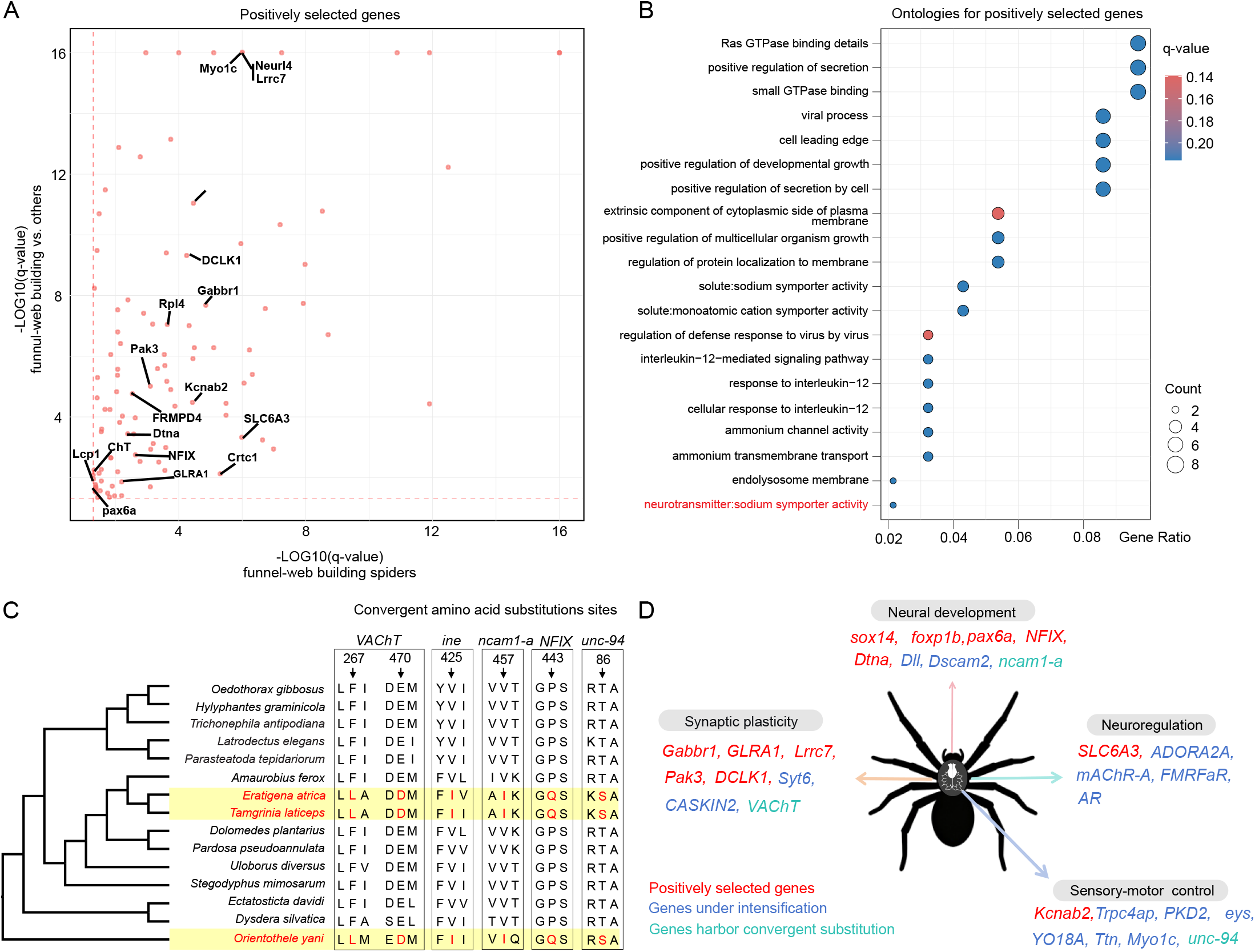
Positive selection, convergent evolution, and integrated molecular basis of funnel-web building behavior. (A) Identification of positively selected genes (PSGs) in sheet-web weaving spiders compared to other spiders. (B) Gene Ontology (GO) enrichment analysis for the positively selected genes. (C) Convergent amino acid substitution sites identified in key neural genes (including *VAChT, ine, ncam1-a, NFIX*, and *unc-94*) across the three funnel-web weaving spiders. (D) A synthesis model showing the enrichment of three evolutionary signals (positive selection, intensified selection, and convergent substitutions) in four core functional categories underlying the funnel-web building phenotype.

In addition, enriched GO categories such as positive regulation of secretion, neurotransmitter:sodium symporter activity, and regulation of protein localization to membrane further support the idea that enhanced neurotransmission efficiency and neuronal connectivity were critical in the evolution of complex funnel-web construction (Figure 4B). Together, these results suggest that positive selection has targeted key neurophysiological pathways facilitating the coordination, learning capacity, and behavioral specialization required for funnel-web architecture.

### Convergent amino acid substitutions associated with funnel-web evolution

To explore potential molecular convergence linked to funnel-web construction, we identified 58 genes showing convergent amino acid substitutions across funnel-web building spiders (Table S6). Among them, five genes (e.g. *VAChT, ine, ncam1-a, NFIX*, and *unc-94*) exhibited highly consistent convergent sites shared exclusively among *T. laticeps, O. yani*, and *E. atrica* (Figure 4C). For example, in *VAChT* (vesicular acetylcholine transporter), a key component of cholinergic synaptic transmission, position 267 displayed a substitution from phenylalanine (F) to leucine (L), and position 470 from methionine (M) to aspartic acid (D) in the three funnel-web spiders, while other spiders retained the ancestral residues. The *ine* gene showed a valine (V) to isoleucine (I) substitution at position 425, and *ncam1-a* displayed a similar change at residue 457. In *NFIX* gene, position 443 was glutamine (G) in funnel-web spiders but proline (P) in other lineages. Similarly, *unc-94* gene showed a substitution from threonine (T) to serine (S) at position 86.

These convergent amino acid changes, particularly within genes involved in synaptic transmission, neuronal differentiation, and cytoskeletal regulation, suggest that adaptive modifications in neural and motor-related pathways may have facilitated the evolution of similar web-building behaviors across distantly related spider lineages.

## Discussion

Spiders exhibit diversity in web architecture and foraging strategies, providing a unique opportunity to investigate the molecular mechanisms underlying behavioral adaptation. Funnel-web spiders inhabit nocturnal, sheltered environments that are rich in prey, and their webs function as mechanosensory traps for ambush predation (Foelix, 2011; Nentwig, 2024). Funnel-web spiders have independently evolved similar funnel-shaped webs with a retreat tube across distinct lineages, notably within the Agelenidae (*T. laticeps, E. atrica*) and Macrothelidae (*O. yani*). This repeated emergence of a complex behavior offers a natural model for studying convergent evolution at both the phenotypic and molecular levels. In this study, we conducted multiple comparative genomics analyses to uncover the molecular basis of funnel-web building behavior, with a particular focus on neural regulatory genes.

Chromosome-level assemblies revealed that *T. laticeps* has 20 chromosomes, *E. atrica* 22, and *O. yani* 46, reflecting notable variation in karyotypic structure. Synteny analyses further demonstrated extensive collinearity between the two Agelenidae species, reflecting their close evolutionary relationship, whereas *O. yani* showed markedly reduced synteny, consistent with its early divergence (∼239 Mya). Such convergence despite deep genomic divergence has been widely recognized in evolutionary biology (Stern, 2013), and even demonstrated at the genomic level in distantly related taxa such as marine mammals (Foote et al., 2015).

Analyses of evolutionary rate shifts revealed contrasting selective pressures across functional gene categories. Genes associated with fundamental cellular maintenance, such as vesicle trafficking, oxidative stress regulation, and Golgi organization, exhibited high evolutionary constraint, reflecting their essential and conserved roles in maintaining cellular homeostasis. In contrast, genes involved in neural signaling, hormone secretion, and developmental regulation evolved more rapidly in funnel-web spiders, suggesting that behavioral innovation may have been facilitated by the accelerated evolution of neural and regulatory pathways. These accelerated changes may enhance sensory processing, motor coordination, and cognitive integration, which likely central to the evolution and maintenance of funnel-shaped web-building behaviors. This pattern, where core cellular processes remain conserved while neural and behavioral genes diversify, is consistent with findings across other animal systems in which complex behaviors have evolved independently (Stern, 2013; Foote et al., 2015). This evolutionary rate heterogeneity suggests that behavioral convergence can occur not through extensive genomic restructuring but via recurrent, fine-scale changes in the tempo of evolution within neural regulatory networks (Reilly et al., 2015).

Funnel-web spiders exhibited both relaxed and intensified selection across distinct gene sets, highlighting the complex interplay between constraint and innovation in behavioral evolution. For example, *FASN*, a key enzyme in lipid biosynthesis that supports neuronal membrane growth and synaptic remodeling (Gonzalez-Bohorquez et al., 2022), showed strong relaxed selection (K = 0.6957, q = 0.0011). Such relaxation may reflect a release of purifying pressure, allowing greater flexibility in metabolic and signaling functions that support behavioral plasticity. In contrast, *Syt6*, a synaptotagmin involved in calcium-dependent neurotransmitter release (Yang et al., 2021), experienced intensified selection (K = 23.3, q = 0.0156), suggesting adaptive fine-tuning of synaptic transmission critical for mechanosensory processing and prey-capture behavior. GO enrichment of relaxed-selection genes indicated shifts in post-transcriptional regulation and innate immune modulation, whereas intensified-selection genes were enriched for pathways related to kinase activity, hormone receptor binding, and neuronal differentiation. Together, these patterns suggest that while some neural genes underwent constraint release to permit adaptive flexibility, others experienced targeted optimization to refine sensory and motor control. This dual selection mode, which integrates the relaxation of regulatory flexibility and the intensification of neural precision, might have collectively promoted the independent emergence of funnel-web building among phylogenetically distant lineages.

Our analyses revealed evidence of positive selection and convergent amino acid substitutions in genes associated with neural signaling and behavioral regulation. Positively selected genes were significantly enriched in pathways related to Ras GTPase binding, vesicle transport, and neurotransmitter:sodium symporter activity, underscoring the importance of neural signaling plasticity in the evolution of web-building strategies. Several candidate genes, including *SLC6A3* (dopamine transporter), *Gabbr1* (GABA-B receptor), and *GLRA1* (glycine receptor), encode key components of neurotransmission systems that regulate sensory perception and motor coordination (Reith et al., 2022; Cediel et al., 2022; Ferraroli et al., 2022). Adaptive modifications in these pathways may have enhanced mechanosensory responses essential for prey detection and web construction. In addition, we identified 58 genes with convergent amino acid substitutions across funnel-web building spiders, indicating shared molecular adaptations underlying similar behaviors. Notably, convergent substitutions occurred in *VAChT, ine, ncam1-a*, and *NFIX*, all of which are involved in neural development, synaptic transmission, or cytoskeletal organization (Prado et al., 2017; Soehnge et al., 1996; Petrovska et al., 2017; Heng et al., 2014). These parallel molecular changes, observed across deeply divergent lineages, highlight how shared selective pressures can drive convergence in neurogenetic pathways, ultimately giving rise to similar behavioral phenotypes. Together, these results support a model in which behavioral convergence emerges from both adaptive fine-tuning of neural function and parallel molecular evolution in synaptic genes. Such integrative signatures of positive selection and amino acid convergence provide compelling evidence that the evolution of funnel-web construction involves repeated exploitation of conserved neurogenetic modules. Furthermore, convergent substitutions were detected in unc-94, a gene encoding tropomodulin, an actin-binding protein critical for sarcomeric organization and muscle function (Stevenson, 2007). This finding implies that, beyond neural regulatory changes, adaptive modifications in cytoskeletal components may also contribute to the behavioral convergence observed in funnel-web spiders.

Our comparative genomic analyses reveal that the convergent evolution of funnel-web construction in spiders is underpinned by coordinated molecular adaptations across multiple neurogenetic modules. To synthesize these findings, we categorized key candidate genes into four interconnected functional groups (Figure 4D): Genes involved in neural development (e.g., *sox14, foxp1b, pax6a, NFIX, Dtna, Dll, Dscam2, ncam1-a*) may influence the formation and wiring of neural circuits that support complex web-building behaviors (Katsuyama et al., 2022; Itoh et al., 2024; Zhang et al., 2010; Heng et al., 2014; Compton et al., 2005; Lah et al., 2014; Petrovska et al., 2017). Genes related to synaptic plasticity (e.g., *Gabbr1, GLRA1, Lrrc7, Pak3, DCLK1, Syt6, CASKIN2, VAChT*) are essential for fine-tuning synaptic transmission and responsiveness to mechanical stimuli transmitted through the web (Cediel et al., 2022; Schaefer et al., 2018; Willim et al., 2024; Boda et al., 2006;

Sossey-Alaoui et al., 1999; Yang et al., 2021; Wang et al., 2024; Prado et al., 2017). Neuroregulatory genes (e.g., *SLC6A3, AR, ADORA2A, mAChR-A, FMRFaR*) potentially modulate neurotransmitter balance and signal propagation, facilitating rapid behavioral responses during prey capture (Miller et al., 2021; Wegener et al., 2022; Rosin et al., 2003; Malloy et al., 2016; Ubuka et al., 2014). Finally, genes associated with sensory-motor control (e.g., *Kcnab2, Trpc4ap, PKD2, eys, YO18A, Ttn, Myo1c, unc-94*) likely contribute to the integration of tactile cues and motor coordination required for web construction and maintenance (Mohamed et al., 2025; Poduslo et al., 2009; Gabrielle et al., 2023; Marques et al., 2025; Buschman et al., 2018; Hanashima et al., 2025; Arif et al., 2019; Stevenson, 2007).

In conclusion, our study thus provides a molecular framework linking genomic signatures to behavioral convergence, illustrating how evolution repeatedly reuses ancient neurogenetic modules to produce similar adaptive outcomes across distant lineages.

## Methods and materials

### Sample collection and preparation

The female spiders of *Tamgrinia laticeps* were collected from Pingwu, Sichuan Province of China, in October 2023. These spiders were subjected to a three-day starvation treatment, cleansed, dissected to remove the abdomen and stomach. Then flash-frozen in liquid nitrogen and stored in a -80°C freezer for subsequent sequencing.

### Sequencing

All the samples were sent to Berry Genomics (Beijing, China) for long-read PacBio HiFi sequencing, Hi-C sequencing and RNA sequencing. Genomic DNA was extracted using the Qiagen Blood & Cell Culture DNA Mini Kit, according to the protocol for PacBio HiFi sequencing. Three 20 kb library were prepared using the PacBio SMRTbell Express Template Prep Kit 2.0 and sequenced on the PacBio Sequel II platform in HiFi mode for three cells. Three Hi-C libraries were constructed through formaldehyde cross-linking, MboI digestion, end-repair, and fragment purification following established methodologies. Short-read sequencing was performed using the Illumina HiSeq PE150 platform. RNA was isolated using TRIzol reagent.The 300 bp librariy was prepared through QiaQuick PCR Kit (Qiagen), then sequenced by Illumina Novaseq 6000.

### Genome assembly and annotation

The genome of *T. laticeps* was assembled using Hifiasm v0.19.5 (Cheng et al., 2021). Hi-C reads were processed with Juicer v1.6.2 (Durand et al., 2021), 3D-DNA v180922 (Dudchenko et al., 2017), and Juicebox to generate a chromosome-level assembly. Potential contaminations were detected and removed using FCS-GX (Astashyn et al., 2024). The completeness and quality of the final assembly were evaluated with BUSCO v5.4.5 ( Waterhouse et al., 2018) using the arachnida_odb10 dataset (n = 2,934).

Repetitive elements were identified using RepeatModeler v2.0.1 (Flynn et al., 2020) and RepeatMasker v4.1.4 (Tarailo-Graovac and Chen, 2009). Non-coding RNAs were annotated with Infernal v1.1.2 (Nawrocki and Eddy, 2013) and tRNAscan-SE v2.0.6 (Chan et al., 2021).

For gene structure annotation, we combined ab initio, transcriptome-based, and protein homology-based evidence using MAKER v3.01.04 (Holt and Yandell, 2011). Ab initio gene predictions were performed with Augustus v3.3.3 (Hoff and Stanke, 2019) and GeneMark-ES/ET/EP v4.48_3.60_lic (Bruna et al., 2020). To improve model accuracy, both gene predictors were initially trained using the BRAKER v3.0.2 pipeline (Bruna et al., 2021). Transcriptome data from the *T. laticeps* were aligned to the genome using HISAT2 v2.2.0 (Kim et al., 2015). The BRAKER pipeline was run with default parameters, and RNA-seq alignments were subsequently assembled into transcripts with StringTie v2.1.3 (Pertea et al., 2015). The resulting transcripts were provided as expressed sequence tag (EST) evidence to MAKER via the “est” option. Protein homology evidence was obtained from the NCBI database, including protein sequences from *Bombyx mori* (GCA_030269925.2) (Lee et al., 2025), *Drosophila melanogaster* (GCA_000001215.4) (Matthews et al., 2015), *Parasteatoda tepidariorum* (GCA_000365465.3) (Zhu et al., 2023), *Stegodyphus mimosarum* (GCA_000611955.2), and *Trichonephila antipodiana* (GigaDB, http://dx.doi.org/10.5524/100868) (Fan et al., 2021).

Gene function annotation was performed using DIAMOND v0.9.24 (Buchfink et al., 2015), InterProScan v5.41–78.0 (Mulder and Apweiler, 2007), and eggNOG-mapper v2.0 (Cantalapiedra et al., 2021). Protein sequences were searched against the UniProtKB/Swiss-Prot database using DIAMOND with the more-sensitive mode, a maximum of one target sequence per query, and an e-value threshold of 1e−5. Functional domains and conserved motifs were identified using InterProScan by querying five databases: Pfam (Mistry et al., 2021), PANTHER (Mi et al., 2016), Gene3D (Lewis et al., 2018), SUPERFAMILY (Pandurangan et al., 2019), and the Conserved Domain Database (CDD) (Marchler-Bauer et al., 2017). Orthology-based functional annotation was further conducted using eggNOG-mapper against the eggNOG v5.0 database (Huerta-Cepas et al., 2019).

In addition, we re-annotated the genomes of *Orientothele yani* (You et al., 2024), *Eratigena atrica* (GCA_965122215.1), *Amaurobius ferox* (GCA_951213105.1) (Schoneberg et al., 2024), *Dolomedes plantarius* (GCA_907164885.2) (Schoneberg et al., 2024) and *Latrodectus elegans* (GCA_030067965.1) (Wang et al., 2022) using the same pipeline.

### Synteny analysis

To investigate genomic collinearity among the three funnel-web spiders *T. laticeps, O. yani*, and *E. atrica*, we performed a genome-wide synteny analysis. Pairwise sequence comparisons were first conducted using BLASTP (Camacho et al., 2009) with an E-value threshold of 1e−5 to identify homologous gene pairs. The resulting BLAST outputs were subsequently analyzed with MCScan (Wang et al., 2012) to detect syntenic blocks across genomes. Finally, the syntenic relationships were visualized using NGenomeSyn (He et al., 2023), enabling clear representation of conserved genomic architectures among species.

### Phylogenetic tree construction and divergence time estimation

To investigate the convergent evolution of funnel-web building behavior in spiders, we performed phylogenomic analyses including *T. laticeps* and several other spider species. The genomes of two additional funnel-web spiders, *O. yani* and *E. atrica*, as well as other representative spiders with diverse web types, were downloaded from public databases. These included the lampshade-web spider *Ectatosticta davidi* (ScienceDB: https://doi.org/10.57760/sciencedb.06872) (Fan et al., 2023), the social bird-nest spider *Stegodyphus mimosarum*, the cribellate orb-weaver *Uloborus diversus* (GCF_026930045.1) (Miller et al., 2022), the orb-weaving spiders *Trichonephila antipodiana, Parasteatoda tepidariorum* (GCA_000365465.3) (Zhu et al., 2023), and *Latrodectus elegans* (GigaDB: http://dx.doi.org/10.5524/102210) (Wang et al., 2022), as well as the ground-dwelling spiders *Dysdera silvatica* (GCA_006491805.2) (Escuer et al., 2022), *Dolomedes plantarius* (GCA_907164875.2) (Schoneberg et al., 2024), and *Pardosa pseudoannulata* (GCA_008065355.1) (Yu et al., 2024).

Shared single-copy orthologous genes were identified from the BUSCO outputs. A total of 2,915 single-copy orthologs were used to construct the phylogenetic tree. Protein sequences were aligned individually using MAFFT v7.471 (Nakamura et al., 2018) with the L-INS-i strategy. The resulting alignments were trimmed with trimAl v1.4 (Capella-Gutiérrez et al., 2009) using the “automated1” method and then concatenated with FASconCAT-G v1.04 (Kück and Longo, 2014).

A maximum-likelihood (ML) tree was inferred using IQ-TREE v2.0.3 (Minh et al., 2020), with model optimization via the rcluster algorithm. Ultrafast bootstrap and Shimodaira–Hasegawa (SH) approximate likelihood ratio tests were performed with 1,000 replicates each, and 10 optimization iterations were applied.

Divergence times were estimated based on fossil calibration points and published references (Magalhaes et al., 2020; Moradmand, Schönhofer, and Jäger, 2014), together with data from the Paleobiology Database (https://paleobiodb.org/) and TimeTree (http://www.timetree.org/). Calibration constraints included the Synspermiata stem (164–175 Mya) and Nephilinae stem (43–47.8 Mya). Divergence time estimation was performed using MCMCTree in PAML v4.10.0 (Yang, 2007), with two independent runs of 50,000 generations each, sampling every five generations and discarding 20% of samples as burn-in.

### Molecular evolution analysis

To explore the molecular basis underlying the convergent evolution of funnel-web construction in spiders, we performed comparative analyses of coding sequence evolution across species. A total of 8,765 shared orthologous genes identified by FastOMA v0.3.4 (Majidian et al., 2025) were used for subsequent analyses.

For each ortholog, the ratio of nonsynonymous to synonymous substitution rates (dN/dS, ω) was estimated with HyPhy (Pond et al., 2005) following the procedure described (Tong et al., 2022). This allowed the assessment of selective constraints acting on funnel-web spiders compared with other web-building lineages. To further examine lineage-specific shifts in selection intensity, RELAX (Wertheim et al., 2015) was applied at the gene level. Genes showing K-values < 1 (q < 0.05) were classified as undergoing relaxed selection, while those with K-values > 1 (q < 0.05) were interpreted as experiencing intensified selection. Multiple testing corrections were performed using the Benjamini– Hochberg approach.

To detect lineage-specific rate shifts associated with funnel-web building spiders, we calculated relative evolutionary rates (RER) using RERconverge v2.6.1 (Kowalczyk et al., 2019). The single-copy ortholog alignments and the species phylogeny described above were used as input, with *T. laticeps, O. yani*, and *E. atrica* defined as focal lineages. For each gene, normalized branch-specific rates were estimated, and the rate parameter (RHO) was used to evaluate acceleration or deceleration in the focal branches. Genes with RHO > 0 (p < 0.05) were considered rapidly evolving genes (REGs), while those with RHO < 0 (p < 0.05) were classified as slowly evolving genes (SEGs).

In addition, evidence of positive selection potentially associated with the evolution of funnel-web traits was detected using BUSTED-PH (Murrell et al., 2015) implemented in HyPhy. Genes exhibiting significant positive selection in funnel-web lineages (q < 0.05 for the foreground branch, q > 0.05 for background branches) were considered to be candidates for adaptive evolution.

### Convergent amino acid substitution analysis

To identify convergent amino acid substitutions (CAAS) associated with funnel-web building behavior, we employed CAAStools v0.6.3 (Barteri et al., 2022). Protein-coding gene alignments derived from single-copy orthologs were used as input. Each alignment was filtered to remove sequences shorter than 50 codons and trimmed to ensure high-quality positional homology.

We defined the three funnel-web building species (*T. laticeps, O. yani*, and *E. atrica*) as the foreground group, and the remaining non-funnel-web spiders as the background group. CAAStools was executed in CAAS detection mode using the default parameters to identify amino acid positions showing convergent substitutions in the foreground group. The resulting convergent sites were statistically evaluated using the simulation module of CAAStools to assess the probability of convergence occurring by chance, and only sites with p < 0.05 were considered significant.

### Functional enrichment and visualization

Genes identified from molecular evolution, convergent amino acid substitution, and evolutionary rate shift analyses were subjected to Gene Ontology (GO) enrichment in R, with correction for gene length bias. Significantly enriched GO terms (FDR < 0.05) were further clustered based on semantic similarity using the simplifyEnrichment R package. Visualization of enrichment results and other data plots in this study were generated using ggplot2.

## Data availability

The data sequenced in this manuscript were submitted in scienceDB dataset.

## Acknowledgements

This work was supported in part by the High Performance Computing(HPC) clusters at Southwest University. This research is supported by the National Natural Science Foundation of China (32570539), the Key Project of Chongqing Municipality (cstc2019jcyj-zdxmX0006), the Science & Technology Fundamental Resources Investigation Program (2024FY100403), and the Fund on survey of Invertebrates from Yintiaoling Nature Reserve (CQS24C00333) by Zhi-Sheng Zhang; and the National Natural Science Foundation of China (32160113), and the Special Program of Science and Technology of Yunnan Province (202002AA100007) by Zi-Zhong Yang.

## Author contributions

Z.Y., C.T., and Z.Z. designed the original research. Z.F. and C.T. performed data analysis and drafted the manuscript. L.W., J.X., L.C., Z.L., and J.K. collected the samples. B.T., T.R., P.W., and W.W. performed formal analyses and visualization of the data. All authors revised and approved the final manuscript.

